# D1/D5 Receptor Activation Promotes Long-Term Potentiation and Synaptic Tagging/Capture in Hippocampal Area CA2

**DOI:** 10.1101/2025.06.01.657216

**Authors:** Kevin Chua, Yee Song Chong, Sreedharan Sajikumar

## Abstract

Hippocampal area CA2 plays an important role in social memory formation. However, CA2 is characterised by plasticity-resistant Schaffer collateral-CA2 (SC-CA2) synapses and highly plastic entorhinal cortex-CA2 (EC-CA2) synapses. Despite abundant dopaminergic input, the relationship between dopamine signalling and area CA2 synaptic plasticity remains unexplored. Here, we show that SKF-38393-mediated Dopamine D1-like receptor (dopamine D_1_ and D_5_ receptors) activation differentially primes CA2 inputs in an NMDAR and protein synthesis-dependent manner. We defined an inverted-U shape relationship between SKF-38393 concentration and EC-CA2 potentiation. Additionally, we observed a priming effect on SC-CA2 plasticity with 50μM SKF-38393, relieving plasticity resistance. We also demonstrated that this effect follows canonical protein kinase A (PKA) signalling. Collectively, our results show that D1R activation primes the CA2 for synaptic plasticity. Thus, we propose a link between neuropsychiatric diseases related to impaired dopamine transmission and deficits in hippocampus-dependent social memory.

## Introduction

The hippocampal area CA2 is involved in the formation of social recognition memory, social aggression, and regulating CA1 excitability ^1–5^. Experimental silencing of the CA2 led to impaired social memory formation. In these experiments, mice were unable to differentiate novel mice from familiar ones ^1,6^. At the circuit level, changes in CA2 firing pattern in response to social stimulation and novel objects were proposed to be a mechanism for how the CA2 encodes the social elements of memory ^2,7^. However, it remains unclear how the CA2 integrates social information with episodic memories.

The CA2 exhibits plasticity resistance at Schaffer Collateral-CA2 synapses (SC-CA2) in the *stratum radiatum* (SR) and robust plasticity at Entorhinal Cortex-CA2 synapses (EC-CA2) in the *stratum lacunosum moleculare* (SLM) ^8,9^. By possessing such unique synaptic properties, the CA2 was proposed to act as a filter to prevent excessive CA1 activity ^8^. For example, post-mortem hippocampal slices of schizophrenic patients exhibited a selective decrease in CA2 inhibitory drive ^10^. This supports accounts of hippocampal hyperactivity in the schizophrenia pathophysiology ^11^. Apart from intrahippocampal projections, the CA2 also receives various neuromodulatory inputs from extrahippocampal structures that modulate synaptic efficacy ^2,9^. These include vasopressinergic projections from the paraventricular nucleus, cholinergic projections from the medial septum and diagonal bands of Broca, and dopaminergic projections from the locus coeruleus (LC) and ventral tegmental area (VTA) ^12–16^. Neuromodulators like vasopressin 1b receptor agonists were shown to promote slow-onset potentiation (SOP) lasting up to 25min in CA2 whole-cell recordings ^17^. However, the effect of dopaminergic modulation on CA2 plasticity remains unclear. The dorsal CA2 robustly expresses Dopamine D1-like receptors (D1R, including dopamine D_1_ and D_5_ receptors) relative to the CA1 and CA3 ^18^. Studies in the CA1 report the involvement of D1R activation in novelty-associated memory, memory consolidation, and spatial memory ^15,16,19–26^. Furthermore, dysfunctions in dopaminergic transmission underlie schizophrenia and autism spectrum disorder ^27–30^. The high density of dopaminergic receptors in CA2, coupled with impaired dopaminergic signalling in schizophrenia, suggests a potential role for CA2 in the pathophysiology of the disorder.

Long-term potentiation (LTP) is a well-regarded correlate for memory formation and is described as a persistent increase in synaptic efficacy ^31,32^. LTP consists of two phases: a short-lasting protein synthesis-independent early phase (early-LTP) and a long-lasting protein synthesis-dependent late phase (late-LTP). In the context of electrically induced LTP, the intensity of tetanic stimulation determines the extent of LTP phase progression, e.g., strong tetanisation (STET) results in the expression of late-LTP. In contrast, weak tetanisation (WTET) results in early-LTP expression. Besides electrical stimulation, neuromodulation differentially facilitates LTP induction in the hippocampus ^24,33–37^. For example, the application of the D1R agonist SKF-38393 with test stimulations resulted in SOP that resembles late-LTP in the CA1 ^24,38,39^. Furthermore, D1R-mediated signalling leads to the synthesis of plasticity-related products (PRPs) ^31,40–42^. PRPs are crucial for both LTP maintenance and the associative properties of LTP. Associativity enables a weakly stimulated pathway – unable to express late-LTP on its own – to do so by accessing resources activated by an independent strongly stimulated pathway in close spatiotemporal proximity. This property is well-illustrated by the synaptic tagging and capture (STC) framework ^43,44^. The STC framework proposes that LTP induction leads to two events: (1) tag setting and (2) PRP synthesis ^31,43,45^. While both WTET and STET are sufficient in synaptic ‘tag’ setting, only STET initiates *de novo* PRP synthesis. These newly synthesized PRPs can be ‘captured’ by ‘tagged’ synapses to transform initially transient LTP into a more persistent form. Since the activity of a synapse can influence another, it suggests that synaptic plasticity itself is malleable, a phenomenon called metaplasticity. Metaplasticity refers to the alteration of the magnitude and length of plasticity due to prior activity-dependent changes in synapses ^46^. Put simply, the LTP induction threshold of synapses can be altered based on prior activity, essentially priming synapses for LTP. This priming effect relies on diverse biochemical mechanisms that create suitable conditions for LTP induction and maintenance.

Newly synthesised PRPs are essential to maintain potentiated synaptic activity. Among the PRPs, Protein Kinase A (PKA) plays an integral role in LTP induction and maintenance, where its downstream targets include Striatum-enriched protein Tyrosine Phosphatase (STEP) and Extracellular signal-regulated Kinase 1/2 (ERK1/2) ^47,48^. In the CA1, ERK1/2 activation via cyclic adenosine monophosphate-protein kinase A (cAMP-PKA) signalling leads to PRP synthesis ^49–51^. STEP is a tyrosine phosphatase highly expressed in the CA2 ^52,53^. As a homeostatic regulator of synaptic plasticity, STEP exerts a tonic brake on LTP induction via ERK1/2 dephosphorylation ^53^. Interestingly, D1R activation in the striatum regulates STEP activity in a cAMP/PKA-dependent manner ^54^. However, it is unclear whether these mechanisms are similar in the CA2.

In this paper, we investigated the effects of SKF-38393-mediated D1R priming in rat hippocampal CA2 via field electrophysiology. Firstly, we illustrated an inverted-U shape dose-response curve of D1R activation on EC-CA2 plasticity. Secondly, we showed that D1R activation primes the SC-CA2 for LTP in a *de novo* protein synthesis- and N-methyl-D-aspartate receptor (NMDAR)-dependent manner. Thirdly, we showed that LTP induction in the SC-CA2 is dependent on EC-CA2 activity. Finally, we showed that SC-CA2 priming involves PKA, STEP, and ERK1/2. Our study points to the role of dopamine signalling in learning and memory, and its possible implications in social-related neuropsychiatric disorders.

## Materials and Methods

### Animals and Preparation of Hippocampal Slices

In this study, a total of 185 hippocampal slices from 59 male Wistar rats (5-7 weeks old) were used for experiments. Wistar rats were obtained from InVivos Pte Ltd (Singapore). All rats were housed in the institutional animal housing facility under 12h light/dark cycle with food and water available *ad libitum*. Animal procedural protocols were approved by the Institutional Animal Care and Use Committee (IACUC) of the National University of Singapore. Rats were briefly anaesthetized with CO_2_, decapitated and the brains were quickly isolated into 2-4°C artificial cerebrospinal fluid (aCSF) containing the following (in mM): 124 NaCl, 2.5 KCl, 2 MgCl_2_•6H_2_O, 2 CaCl_2_•2H_2_O, 1.25 NaH_2_PO_4_, 26 NaHCO_3_ and 17 D-Glucose. These concentrations were adapted and modified from Chevaleyre and Siegelbaum^8^. The aCSF pH was kept between pH 7.2-7.4 and saturated with carbogen (95% O_2_ and 5% CO_2_). Right hippocampi were isolated and sliced transversely into 400μM slices via a manual chopper (Stoelting, Wood Dale, Illinois). These transverse hippocampal slices were laid onto a nylon net in an interface chamber (Scientific Systems Design) and incubated at 32±1°C (PTC03, Scientific System Design Inc., Canada) with an aCSF flow rate of 1ml/min and constant carbogen bubbling for at least 3-4h before any experiment.

### Field Potential Recordings

For two-pathway experiments, three monopolar lacquer-coated stainless-steel electrodes (5 MΩ; AM-Systems, Sequim, Washington) were placed in the CA2. Among the three electrodes, two were utilised as stimulating electrodes, one positioned in the CA2 SR layer to stimulate CA3➔CA2 fibres (SC-CA2) and the other positioned in the CA2 SLM layer to stimulate EC➔CA2 fibres (EC-CA2) (Figure 1a). Field excitatory postsynaptic potentials (fEPSPs) derived from stimulation of the EC-CA2 and SC-CA2 inputs were recorded from the CA2 dendritic region between these two inputs via the recording electrode (Figure 1a). The recordings were amplified via a differential amplifier (Model 1700, A-M Systems) and digitized via an analog-to-digital converter (Power1401, Cambridge Electronic Design). The initial slope value (mV/ms) of fEPSPs was recorded and displayed through the custom-developed Intracell (IfN) software. A paired-pulse facilitation protocol was conducted to test the pathway independence of the two inputs ^55,56^. After a 3-4h preincubation period for slice recovery, an input-output curve was generated (afferent stimulation intensity against fEPSP slope value) for each stimulated input. The test stimulation strength was determined based on the fEPSP response at 40% of the maximal fEPSP slope value. A stable baseline was recorded for 30min at this intensity before any pharmacological intervention or tetanic stimulation. Four sweeps of 0.2Hz biphasic, constant current pulses (pulse duration of 0.1ms) were given every 5min for baseline and post-induction recording. The average slope value from the four sweeps was considered as one point for the fEPSP (%) against time. A weak tetanisation (WTET) protocol comprising of one high-frequency stimulation (HFS) at 100Hz, 21 biphasic current pulses, biphasic burst with a 0.2ms pulse duration. This protocol was shown to be able to induce a transient and protein synthesis-independent form of LTP that typically lasts for around 2-3h before returning to baseline ^42^. This protocol was used for all SC-CA2 WTET induction. A strong tetanisation (STET) protocol comprising three trains (10-min intervals) of 100Hz, 100 pulses, biphasic burst with 0.2ms pulse duration. In the CA1, pathway independence for D1R-mediated SOP could be achieved by selectively silencing test stimulations to a pathway ^24,39,40^. Therefore, for the silenced EC-CA2 experiment, a stable baseline of EC-CA2 was recorded before test stimulations were silenced. The stimulations were turned back on only after 30min post-WTET to prevent EC-CA2 SOP. This was to ensure minimal influence from EC-CA2 onto SC-CA2 D1R priming.

**Figure 1.**
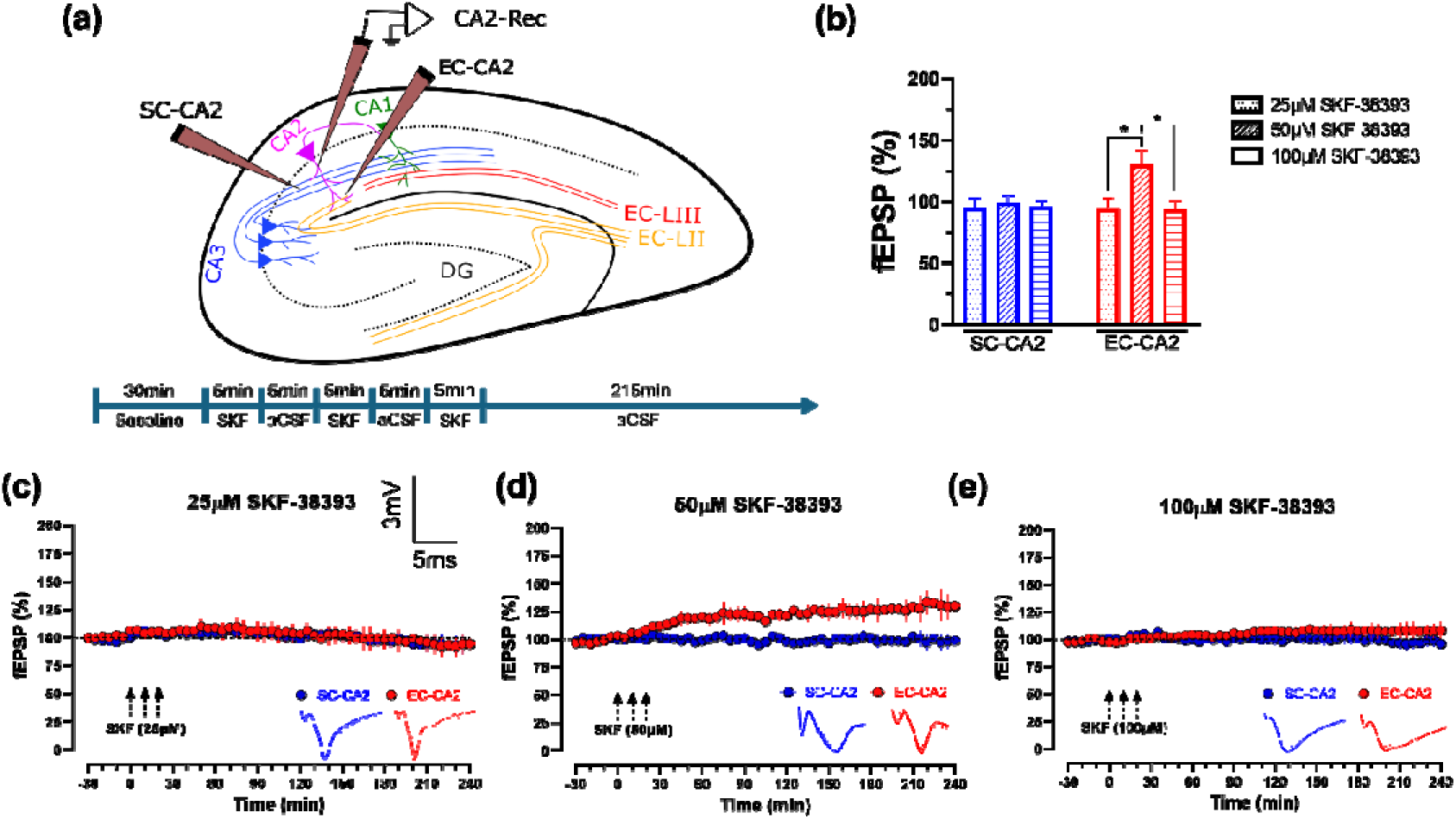
50μM SKF-38393 induces SOP in EC-CA2. (a) (Top) Schematic representation of a hippocampal slice showing the location of electrodes in the CA2 region. Recording electrode was positioned within the CA2 dendritic region. The stimulating electrodes flanked the recording electrode in the SR (SC-CA2) and SLM (EC-CA2). (Bottom) Timeline for the SKF-38393 protocol. (b) A histogram of mean fEPSP percentage in EC-CA2 and SC-CA2 for 25μM, 50μM, 100μM SKF-38393 application at the 240^th^ min. When all three concentrations were compared together, there was a significant difference in EC-CA2 fEPSP percentage at the 240^th^ min (Kruskal-Wallis test, *P*=0.0029). Dunnett’s post hoc test showed no significant difference between 25μM and 100μM SKF-38393 groups (*P*>0.9999) but a significant difference between 25μM and 50μM (*P*=0.0332), and 100μM and 50μM (*P*=0.0150). No significant change in SC-CA2 fEPSP was observed. Asterisks indicate significant differences between groups (Kruskal-Wallis test, \**P*<0.05, \*\**P*<0.01, \*\*\**P*<0.001). Error bars indicate ±SEM. (c) Application of 25μM SKF-38393 did not result in significant potentiation in EC-CA2 (red circles, n=6). (d) Application of 50μM SKF-38393 did not result in significant potentiation in EC-CA2 (red circles, n=7). (e) Application of 100μM SKF-38393 did not result in significant potentiation in EC-CA2 (red circles, n=6). Three dotted arrows represent SKF-38393 5-min application intervals. Scale bars for all the traces vertical: 3mV; horizontal: 5ms.

### Pharmacology

D1R partial agonist SKF-38393 (#D047, Sigma-Aldrich) was prepared as a 50mM stock diluted in 0.1% DMSO and stored at -20°C. DMSO at 0.1% does not affect basal transmission and does not affect control recordings ^57^. Before drug application, the stock solution was diluted to a final concentration of either 25^μ^M, 50_μ_M or 100_μ_M in aCSF. SKF-38393 was bath applied to the interface chamber at ∼1ml/min for 5min with a 5-min interval of aCSF perfusion (Figure 1a). This paradigm was suggested to emulate spaced HFS-like LTP induction in hippocampal CA1 neurons ^39,58^. The protein synthesis inhibitor emetine dihydrochloride (#E2375, Sigma-Aldrich) was stored as a 20mM stock prepared in de-ionised water and diluted to a 20_μ_M working solution in aCSF before application. NMDAR antagonist D-AP5 (#A8054, Sigma-Aldrich) was stored as a 50mM stock prepared in de-ionised water and diluted to a 50_μ_M working solution in aCSF before application. Based on the experiment, either Emetine or AP5 was bath applied to the interface chamber for a total of 60min after baseline. 30min into the drug application, 50_μ_M SKF-38393-drug mixture was used as in the protocol mentioned above. However, instead of aCSF perfusion during the spaced interval, it was substituted with either Emetine- or D-AP5-aCSF respectively.

All concentrated stocks were made freshly every week before experimentation and used within the week.

### Western Blot

Control and SKF-38393-treated slices were flash-frozen in liquid nitrogen and stored at - 80°C until use. The CA2 was apportioned under a microscope from these flash-frozen slices. Each sample contained 8 slices. Total protein was extracted using T-PER Tissue Protein Extraction Kit (Prod#78510, Thermo Fischer Scientific Inc., USA) and HALT^TM^ Protease Inhibitor Cocktail Kit (Prod#78840, Thermo Fischer Scientific Inc., USA). A colourimetric assay dye concentrate was used to quantify protein levels in samples (Quick Start Bradford Protein Assay, 5000205, Bio-Rad, CA., USA). 20μg of protein extracts were separated in 10% SDS-polyacrylamide gels and wet transferred to polyvinylidene difluoride (PVDF) transfer membranes. The membranes were blocked for 1h with 5% bovine serum albumin (A2153, Sigma-Aldrich, MO, USA) and subsequently incubated with the respective primary antibodies overnight at 4°C. The primary antibodies used were rabbit anti-phosphorylated-PKA C (Thr197) (1:1 000, #4781, Cell Signaling Technology, USA), rabbit anti-PKA Cα (1:1 000, #4782, Cell Signaling Technology, USA), rabbit anti-phosphorylated-p44/42 MAPK Thr202/Tyr204 (1:1 000, #9101, Cell Signaling Technology, USA), rabbit anti-p44/42 MAPK (1:1 000, #9102, Cell Signaling Technology, USA), mouse anti-STEP (1:200, sc-23892, Santa Cruz Biotechnology), and mouse anti-GAPDH (1:3 000, #97166, Cell Signaling Technology, USA), and incubated with their respective secondary antibodies conjugated with horseradish peroxidase (rabbit and mouse, Cell Signaling Technology, USA). The protein bands were detected via chemiluminescence (SuperSignal West Pico PLUS Chemiluminescent Substrate Kit, Thermo Scientific, USA) and imaged via a chemiluminescence imaging machine (Azure 400, Azure Biosystems, USA). ImageJ was utilised to quantify the protein bands ^59^, with bands normalised to their corresponding total protein or GAPDH.

### Statistical Analysis

The fEPSP slope values per time point were expressed as a percentage against the average 30min baseline values in all experiments. GraphPad Prism 10.0 was used to plot graphs and perform statistical analyses on the average time-matched, normalised data across replicated experiments. The fEPSP graphs are plotted against time as ‘mean ± standard error of mean (SEM)’. All experiments were subjected to non-parametric tests as normality cannot be assumed at small sample sizes. Wilcoxon signed-rank test was used when comparisons were made within a group. Mann-Whitney U test was used to compare values between groups. For multiple comparisons between three or more groups, the Kruskal-Wallis test was used. Dunnett’s post hoc test was used to determine differences between control and experimental groups. Unpaired t-test was used to analyse western blot results. The statistical significance was assumed from *P*<0.05 (*P*<0.05: *, *P*<0.01: **, *P*<0.001: ***), and the number of slices used were denoted as ‘n’. Every experimental group consists of a minimum of six biological variants.

## Results

### 50μM **SKF-38393 induces slow-onset potentiation in EC-CA2 but not SC-CA2**

Since most studies in the CA1 use a range between 50_μ_M to 100_μ_M SKF-38393 ^24,38,39,50,60^, we decided to use 50_μ_M as the midpoint concentration to gauge the response of the CA2 between lower (25_μ_M) and higher (100_μ_M) concentrations. There was no significant potentiation observed in EC-CA2 and SC-CA2 from the 25_μ_M and 100_μ_M groups (Figure 1c & 1e). In contrast, 50μM SKF-38393 induced SOP in the EC-CA2 that lasted throughout the recording (Figure 1d, 130.1±11.5% vs 97.9±1.2% (baseline), Wilcoxon test, *P*=0.0156, n=7). The Kruskal-Wallis test revealed a significant difference between the 50μM SKF-38393 EC-CA2 endpoint (240min) mean fEPSP slope value and the EC-CA2 endpoints of 25μM and 100μM (Figure 1b, *H*=9.844, *P*=0.0029). Dunnett’s post hoc test revealed no significant difference between 25μM and 100μM groups (*P*>0.9999) but a significant difference between 25μM and 50μM (*P*=0.0332), and 100μM and 50μM (*P*=0.0150). There were no significant differences in SC-CA2 baseline and endpoint potentiation for 50μM SKF-38393 (98.8±6.4% vs 100.8±1.7% (baseline), Wilcoxon test, *P*=0.5781, n=7). Furthermore, the Kruskal-Wallis test revealed no significant difference between SC-CA2 endpoint potentiations across all concentrations (Figure 1b, *H*=9.844, *P*=0.6008). Since SOP was not observed in the SC-CA2 upon SKF-38393 application, it affirms its plasticity-resistant nature.

These observations show that 50μM SKF-38393 sufficiently induces SOP resembling late-LTP in EC-CA2. As such, this concentration was used for the remainder of the study.

### D1R activation primes the SC-CA2 for late-LTP

Since SKF-38393 did not induce SOP in the SC-CA2, we wanted to determine if the SC-CA2 was primed by D1R activation. A WTET on its own induced a transient potentiation that returned to baseline level within minutes (Figure 2a, 101.9±2.9% vs 99.7±0.5% (baseline), Wilcoxon test, *P*=0.5469, n=8). When the WTET was delivered to the SC-CA2 synapse after SKF-38393 bath application, a priming effect occurred as there was a significant difference between SC-CA2 endpoint potentiation compared to its baseline (Figure 2b, 140.7±8.1% vs 101.7±1.9% (baseline), Wilcoxon test, *P*=0.0156, n=7). Also, EC-CA2 potentiation at the end of 240min was significantly different from the baseline (Figure 2b, 120.4±7.6% vs 99.7±1.6% (baseline), Wilcoxon test, *P*=0.0313, n=7). Additionally, there was a significant difference when endpoint potentiations were compared between untreated WTET SC and SKF+WTET SC (Figure 2c, Mann-Whitney U test, *U*=7, *P*=0.0140). This suggests that D1R activation had primed SC-CA2 synapses, allowing WTET to induce late-LTP, when it would normally induce a transient potentiation.

**Figure 2.**
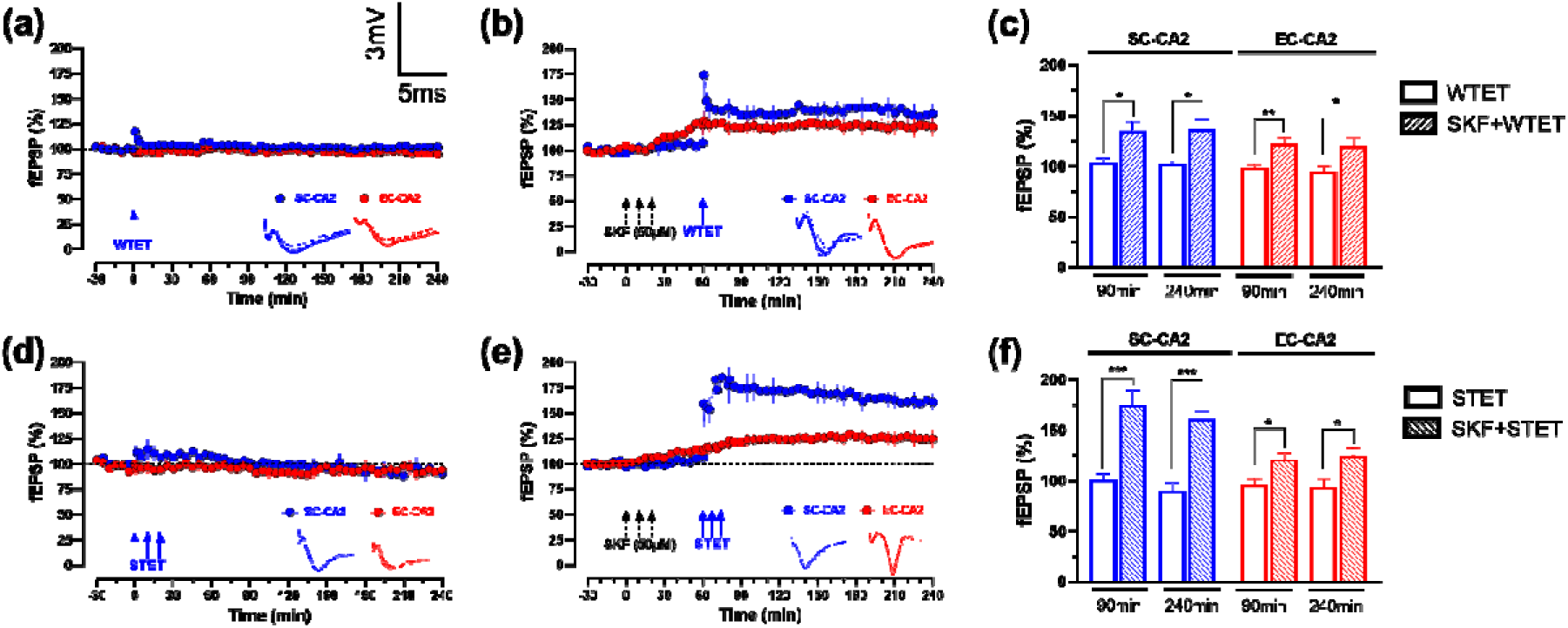
D1R activation via 50μM SKF-38393 primes SC-CA2 for LTP. (a) Delivery of WTET to untreated control SC-CA2 resulted in a transient spike in potentiation that quickly decays back to baseline (blue circles, n=8). (b) Delivery of WTET to 50μM SKF-38393-treated SC-CA2 resulted in late-LTP (blue circles, n=7). SOP was observed in EC-CA2 that lasted until the end of the recording (red circles, n=7). (c) A histogram of mean fEPSP percentage in EC-CA2 and SC-CA2 between untreated WTET control and SKF+WTET SC at the 90^th^ and 240^th^ min. There was a significant difference in EC-CA2 potentiation between untreated WTET and SKF+WTET SC at 90^th^ min (*P*=0.0013) and 240^th^ min (*P*=0.0313). There was a significant difference in SC-CA2 potentiation between untreated WTET and SKF+WTET SC at 90^th^ min (*P*=0.0205) and 240^th^ min (*P*=0.0156). (d) Delivery of STET to untreated control SC-CA2 resulted in a transient spike in potentiation that quickly decays back to baseline (blue circles, n=7). (e) Delivery of STET to 50μM SKF-38393-treated SC-CA2 resulted in late-LTP (blue circles, n=9). SOP was observed in EC-CA2 that lasted until the end of the recording (red circles, n=9). (f) A histogram of mean fEPSP percentage in EC-CA2 and SC-CA2 between untreated STET control and SKF+STET SC at the 240^th^ min. There was a significant difference in EC-CA2 potentiation between untreated STET and SKF+STET SC at 90^th^ min (*P*=0.0229) and 240^th^ min (*P*=0.0469). There was a significant difference in SC-CA2 potentiation between untreated STET and SKF+STET SC at 90^th^ min (*P*=0.0007) and 240^th^ min (*P*=0.0012). Asterisks indicate significant differences between untreated and SKF-treated groups at the 90^th^ min or 240^th^ min (Mann-Whitney U test, \**P*<0.05, \*\**P*<0.01, \*\*\**P*<0.001). One solid arrow represents WTET for the induction of early[LTP. Three solid arrows represent STET for the induction of late[LTP. Error bars indicate ±SEM. Scale bars for all the traces vertical: 3mV; horizontal: 5ms.

Similar to WTET, delivery of STET only induced transient potentiation (Figure 2d, 90.0±8.1% vs 98.5±2.5% (baseline), Wilcoxon test, *P*=0.2188, n=9) in untreated SC-CA2. When STET was delivered to the SKF-38393 bath-applied SC-CA2 synapses, it potentiated with the endpoint significantly different from baseline (Figure 2e, 163.0±9.3% vs 99.4±1.2% (baseline), Wilcoxon test, *P*=0.0156, n=9). Also, EC-CA2 potentiation at the end of 240min was significantly different from baseline (Figure 2e, 114.8±5.7% vs 98.9±1.4% (baseline), Wilcoxon test, *P*=0.0469, n=9). Additionally, there was a significant difference when endpoint potentiations were compared between untreated STET SC and SKF+STET SC (Figure 2f, Mann-Whitney U test, *U*=1, *P*=0.0012). This reaffirms that plasticity resistance was lifted upon D1R priming.

To investigate if D1R priming enhanced LTP induction, baseline potentiation was compared to the 90^th^ min. There were significant differences in SC-CA2 potentiation between 90^th^ min vs baseline in SKF+WTET SC (135.0±9.2% vs 101.6±1.9% (baseline), Wilcoxon test, *P*=0.0313, n=7) and SKF+STET SC (174.5±15.2% vs 97.9±1.9% (baseline), Wilcoxon test, *P*=0.0039, n=9). Similarly, a significant difference was found in EC-CA2 potentiation between 90^th^ min vs baseline potentiation for SKF+WTET SC (122.3±6.5% vs 100.3±1.8% (baseline), Wilcoxon test, *P*=0.0313, n=7) and SKF+STET SC (121.0±6.5% 98.6±1.6% (baseline), Wilcoxon test, *P*=0.0039, n=9). Furthermore, significant differences were found in SC-CA2 potentiation between WTET SC vs SKF+WTET SC (Figure 2c, 103.4±4.2% vs 135.0±9.2%, Mann-Whitney U test, *U*=8, *P*=0.0205) and STET SC vs SKF+STET SC (Figure 2f, 100.9±6.3% vs 174.5±15.3%, Mann-Whitney U test, *U*=2, *P*=0.0007). Similarly, significant differences were found in EC-CA2 90^th^ min potentiation between WTET SC vs SKF+WTET SC (Figure 2c, 98.4±2.8% vs 122.3±6.5%, Mann-Whitney U test, *U*=7, *P*=0.0013) and STET SC vs SKF+STET SC (Figure 2f, 95.7±6.2% vs 121.3±6.5%, Mann-Whitney U test, *U*=10, *P*=0.0229). Together, these results imply that D1R priming with SKF-38393 reduced the LTP induction threshold in SC-CA2. Regarding LTP maintenance, there was no significant difference in endpoint potentiation between SC-CA2 from the SKF+WTET SC and SKF+STET SC (136.2±10.1% vs 161.1±7.7%, Mann-Whitney U test, *U*=17, *P*=0.1416). Perhaps the priming effect of SKF-38393 was not strong enough such that different HFS intensities produced no difference in the potentiation magnitude of LTP maintenance. Nevertheless, these results suggest that both EC- and SC-CA2 benefitted from D1R priming.

### D1R priming of the CA2 is dependent on EC-CA2 activity, *de novo* protein synthesis, and NMDAR activation

The induction and maintenance of LTP requires the co-activation of dopaminergic and glutamatergic receptors in the CA1 ^39,40,61^. We reasoned that D1R agonist application is similar to the strong tetanisation of synapses ^38,39^. Therefore, to achieve input specificity between EC-CA2 and SC-CA2, we selectively silenced the EC-CA2 during SKF-38393 bath application, 60min before and after WTET SC-CA2, for a total of 120min ^24^. This distinguishes whether the SC-CA2 can recruit its own PRPs or utilise PRPs from activated EC-CA2. As shown in Figure 3b, there was no statistical difference between baseline and endpoint potentiation for both EC-CA2 (Figure 3b, 94.6±7.3% vs 99.9±1.4% (baseline), Wilcoxon test, *P*=0.6953, n=11) and SC-CA2 (Figure 3b, 104.5±6.1% vs 98.8±0.5% (baseline), Wilcoxon test, *P*=0.4258, n=11). Between the SC-CA2 synapses of SKF+WTET and Silenced EC SKF+WTET groups, there were significant differences during the induction (Figure 3e, 174.2±13.8% vs 116.3±5.4%, Mann-Whitney U test, *U*=3, *P*=0.0004) and maintenance (Figure 3e, 136.2±10.1% vs 104.5±6.1%, Mann-Whitney U test, *U*=13, *P*=0.0204) phases. These observations suggest that SC-CA2 LTP induction requires EC-CA2 activity such as PRP synthesis, on top of D1R activation. However, the presence of a trend in SC-CA2 potentiation suggests that there may be some priming activity that is just not statistically significant (Figure 3b).

**Figure 3.**
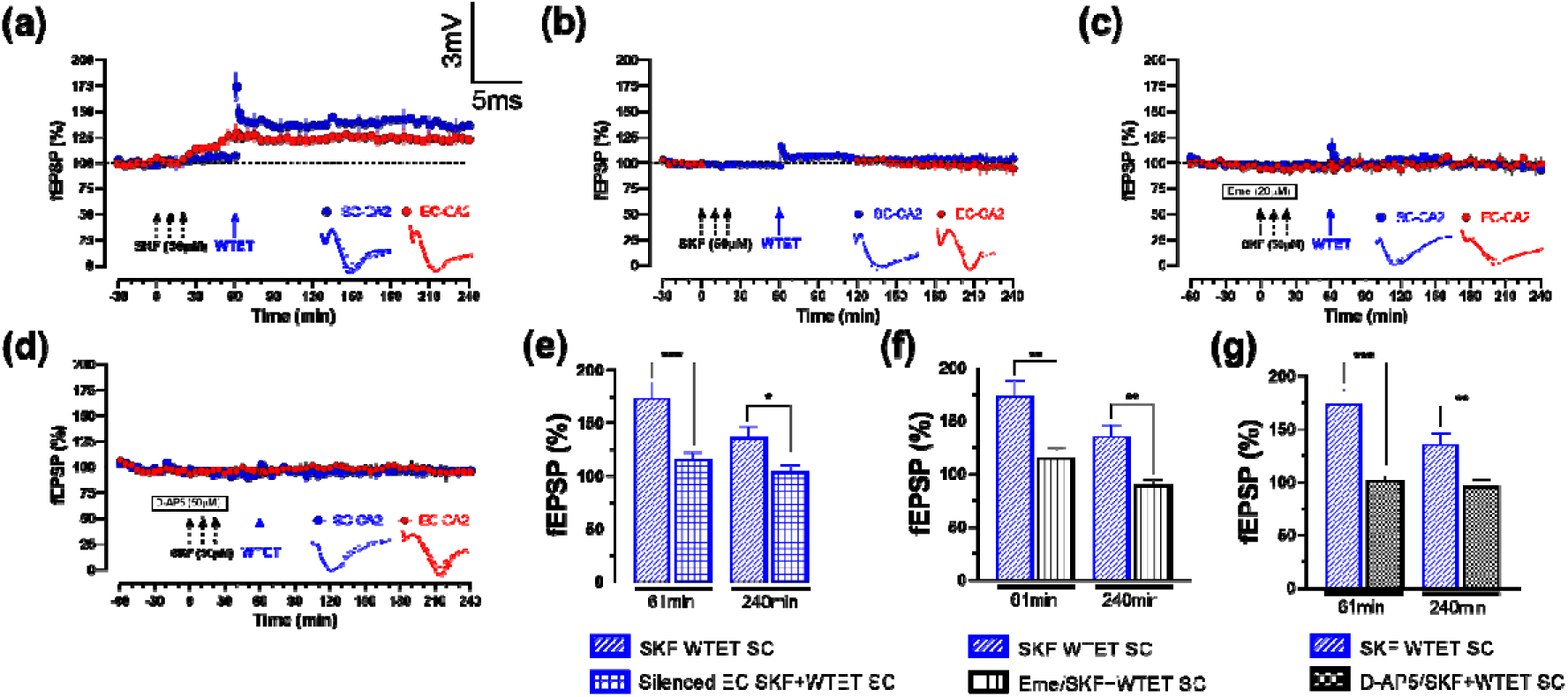
D1R priming effect in SC-CA2 requires EC-CA2 activity, *de novo* protein synthesis and NMDAR activation. (a) Delivery of WTET to 50μM SKF-38393-treated SC-CA2 resulted in late-LTP (blue circles, n=7). SOP was observed in EC-CA2 that lasted until the end of the recording (red circles, n=7). (b) Delivery of WTET to 50μM SKF-38393-treated SC-CA2 did not result in LTP when EC-CA2 was silenced during SKF-38393 bath application and WTET (blue circles, n=11). (c) Delivery of WTET to 50μM SKF-38393-treated SC-CA2 did not result in LTP when protein synthesis inhibitor Emetine (Eme) was applied before and during SKF-38393 bath application (blue circles, n=8). (d) Delivery of WTET to 50μM SKF-38393-treated SC-CA2 did not result in LTP when NMDAR inhibitor AP-5 was applied before and during SKF-38393 bath application (blue circles, n=8). (e-g) Histograms of mean fEPSP percentage in SC-CA2 between SKF+WTET SC and either Silenced EC SKF+WTET SC, Eme/SKF+WTET SC or D-AP5/SKF+WTET SC at the point of induction (61^st^ min) and maintenance (240^th^ min). (e) There was a significant decrease in potentiation between SKF+WTET SC and Silenced EC SKF+WTET SC during induction (*P*=0.0004) and maintenance (*P*=0.0204). (f) There was a significant decrease in potentiation between SKF+WTET SC and Eme/SKF+WTET SC during induction (*P*=0.0041) and maintenance (*P*=0.0022). (g) There was a significant decrease in potentiation between SKF+WTET SC and D-AP5 SKF+WTET SC during induction (*P*=0.0003) and maintenance (*P*=0.0093). Asterisks indicate significant differences between groups (Mann-Whitney U test, \**P*<0.05, \*\**P*<0.01, \*\*\**P*<0.001). One solid arrow represents WTET for the induction of early[LTP. Error bars indicate ±SEM. Scale bars for all the traces vertical: 3mV; horizontal: 5ms.

To investigate whether D1R priming is dependent on *de novo* protein synthesis, protein synthesis inhibitor Emetine was applied for 60min after baseline and during SKF-38393 application. There was no statistical difference between endpoint potentiation and baseline for EC-CA2 (Figure 3c, 97.2±7.0% vs 98.77±0.9% (baseline), Wilcoxon test, *P*=0.7422, n=8) and SC-CA2 (Figure 3c, 90.0±5.2% vs 98.7±1.8% (baseline), Wilcoxon test, *P*=0.2500, n=8), thus implying that D1R priming depends on *de novo* protein synthesis. Furthermore, there were significant differences during the induction (Figure 3f, 174±13.8% vs 115.7±8.3%, Mann-Whitney U test, *U*=3, *P*=0.0041) and maintenance (Figure 3f, 136.2±10.1% vs 89.8±5.2%, Mann-Whitney U test, *U*=3, *P*=0.0022) phases compared to SKF+WTET SC. Together, it shows that (1) D1R priming is protein synthesis-dependent and (2) D1R-primed SC-CA2 likely recruits PRPs but alone is inadequate for LTP.

To investigate whether D1R priming of CA2 is NMDAR-dependent, 50μM NMDAR antagonist D-AP-5 was co-applied with SKF-38393. Similar observations were made that there was no statistical difference between endpoint potentiation and baseline of EC-CA2 (Figure 3d, 95.2±8.4% vs 97.3±1.2% (baseline), Wilcoxon test, *P*=0.9453, n=8) and SC-CA2 (Figure 3d, 96.7±5.9% vs 99.8±1.8% (baseline), Wilcoxon test, *P*=0.6406, n=8). There were significant differences during the induction (Figure 3g, 174.2±13.4% vs 102.1±3.2%, Mann-Whitney U test, *U*=0, *P*=0.0003) and maintenance (Figure 3g, 136.2±10.1% vs 96.69±5.9%, Mann-Whitney U test, *U*=6, *P*=0.0093) phases compared to SKF+WTET SC. As such, D1R priming of the CA2 is indeed NMDAR-dependent, consistent with previous reports in the CA1 ^38–40^.

To conclude, these observations show that (1) D1R activation primes the SC-CA2, (2) this priming effect is dependent on both D1R activation and EC-CA2 activity, and (3) also protein synthesis- and NMDAR-dependent.

### D1R priming in SC-CA2 follows canonical PKA signalling and attenuates STEP activity

Similar to the CA1, D1R and NMDAR co-activation results in SOP in the CA2 (results shown here). However, it is unknown whether D1R activation in the CA2 follows the canonical PKA signalling cascade. As such, we asked whether the CA2 adheres to this canonical signalling cascade after D1R activation. We have shown in an earlier study that D1R activation via SKF-38393 in the CA1 induces ERK1/2-dependent SOP in a dose-dependent manner ^24^. To investigate the mechanism of priming the SC-CA2, only the SC-CA2 was test stimulated for untreated and SKF-38393-treated slices. We made the following assumptions, (1) any intracellular activity related to priming the SC-CA2 is localised to the test stimulation site, and (2) without test stimulation, the EC-CA2 will not induce PRPs required for SOP as we have shown in the Silenced EC experiment. Indeed, D1R priming via SKF-38393 in the area CA2 follows canonical PKA signalling via the upregulation of p-PKA (Figure 4b, 1.8±0.2 vs 0.9±0.1 (control), *P*=0.0022) and p-ERK1/2 (Figure 4d, 1.5±0.2 vs 1.0±0.01 (control), *P*=0.0433), and attenuation of STEP (Figure 4f, 0.7±0.1 vs 1.0±0.003 (control), *P*=0.0010).

**Figure 4.**
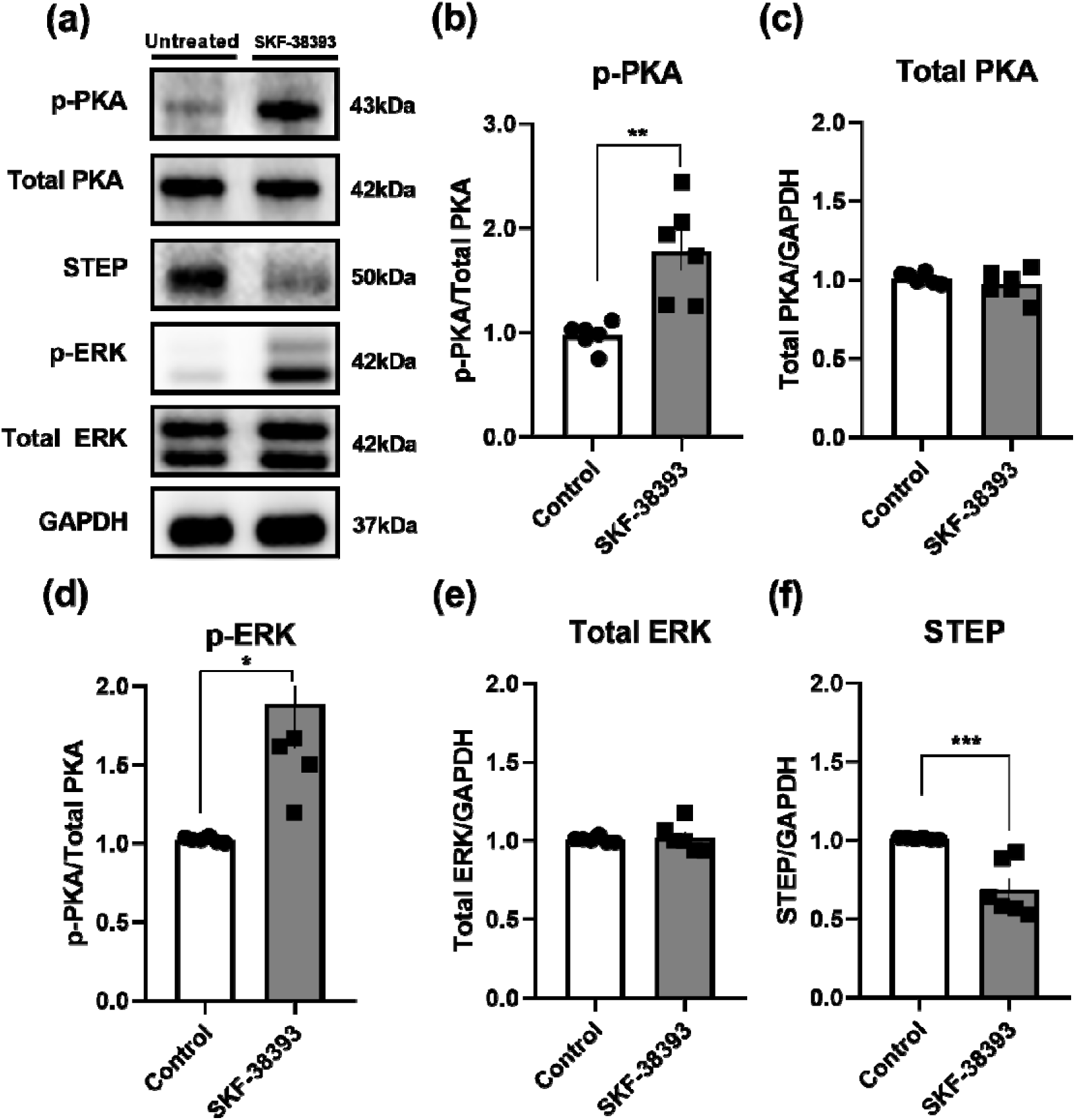
D1R priming effect upregulates PKA and ERK activity while downregulates STEP activity. (a) Western blot analysis of p-PKA, Total PKA, p-ERK, Total ERK and STEP levels between untreated control and SKF-38393-treated groups (n=6 for all groups). (b) p-PKA was significantly increased in the SKF-38393-treated group compared to the untreated group (*P*=0.0022). (c) There was no significant difference between total PKA levels (*P*=0.3528). (d) p-ERK was significantly increased in the SKF-38393-treated group compared to the untreated group (*P*=0.0433). (e) There was no significant difference between total ERK levels (*P*=0.7621). (f) STEP was significantly decreased in the SKF-38393-treated group compared to the untreated group (*P*=0.0010). (b-e) The values of the individual groups were calculated in relation to the control group while GAPDH serves as a loading control. Asterisks indicate significant differences between groups (unpaired t-test, \**P*<0.05, \*\**P*<0.01, \*\*\**P*<0.001). Error bars indicate ±SEM.

## Discussion

Dopaminergic modulation is critical for learning and memory ^37,39^. Numerous works investigating the effect of dopaminergic signalling in the hippocampus regarding memory consolidation and synaptic plasticity have almost been exclusive to the area CA1 and dentate gyrus ^19,22,24,38,62–66^. However, the effects of D1R-mediated plasticity in the area CA2 have not been characterised until now. Our study provides the first compelling evidence that D1R priming facilitates LTP in the area CA2. Furthermore, this effect involves PKA signalling and is dependent on *de novo* protein synthesis and NMDAR activation. We also show that this effect of priming is dependent on the dynamics between EC-CA2 and SC-CA2 activity.

We demonstrated for the first time that the relationship between D1R agonist SKF-38393 concentration and the magnitude of potentiation in the EC-CA2 can be described in an inverted U-shape dose-response curve as only 50μM SKF-38393 was seen to produce significant SOP. 25μM SKF-38393 was likely insufficient to induce SOP in EC-CA2, an effect similarly observed in the CA1 ^24^. While the reason for the lack of SOP in 100μM SKF-38393 remains unclear, future work could explore possible negative feedback mechanisms in the CA2. In contrast, SC-CA2 did not potentiate across all concentrations. Instead, D1R activation primes the SC-CA2 for persistent LTP expression in response to WTET or STET, which normally only elicits transient potentiation as seen in the untreated groups. This implies that D1R activation triggers a metaplastic shift in the CA2 LTP threshold within a ‘goldilocks’ level of dopaminergic stimulation. In contrast, SKF-38393 bath application to the CA1 exhibits a linear dose-response relationship ^24^. This reiterates the intrinsic differences in plasticity expression between the CA1 and CA2 and supports the notion that the CA2 is an information filter, only allowing specific social-related information to be encoded ^67,68^. Various studies suggest that CA2 plasticity is heavily influenced by robust feedforward inhibition from the CA3 and the pro-inhibitory CA2 milieu – e.g., dense perineuronal nets and parvalbumin-positive interneuron population, and high calcium ion buffering and extrusion ^8,67,69–73^. As such, a myriad of internal and external factors regulate CA2 plasticity ^67,74^. Interestingly, chronic silencing of CA2 transmission results in hippocampal hyperexcitability. These findings poise the CA2 as a key regulator of hippocampal excitability by balancing excitatory and inhibitory dynamics ^8,73,75^. Since this property is observed in normal physiological states, it suggests that plasticity suppression in the CA2 has functional importance, essentially responding to the right signals at the right time. This is congruent with our findings that SC-CA2 plasticity is dependent on D1R-mediated EC-CA2 activity as silencing the EC-CA2 during SKF-38393 application abolished LTP in SC-CA2. It implies that the effects of D1R activation at SC-CA2 synapses alone were insufficient to initiate adequate PRP synthesis for capture, resulting in the loss of the phenotype. Together, these observations suggest that the CA2 is selective in processing information.

Regulator of G protein Signaling (RGS14) is commonly regarded as a major suppressive factor of plasticity selectively enriched in the CA2 ^67^. However, RGS14’s inhibitory effect on LTP is independent of G_i/o_-cAMP signalling ^76^. Instead, extensive RGS14-mediated calcium extrusion and buffering restrict CA2 plasticity ^67^. Similarly, Purkinje Cell Protein 4 (PCP4), a calmodulin modulator selectively enriched in the CA2, was proposed to limit plasticity by regulating calcium extrusion via plasma membrane ATPases ^2^. Collectively, multiple plasticity-suppressing proteins and the pro-inhibitory CA2 milieu strictly regulate CA2 plasticity ^2,75,77^. However, as we were interested in the effects of dopaminergic signalling on CA2 plasticity, we looked at mediators of plasticity downstream of D1R signalling. D1Rs are G protein-coupled receptors widely expressed in rat hippocampal CA2 ^78,79^. Activation of D1Rs upregulates PKA activity in a cAMP-dependent manner ^61,66,80,81^. PKA activation upregulates α-amino-3-hydroxy-5-methyl-4-isoxazolepropionic acid receptor (AMPAR) synthesis, trafficking, insertion, and conductance, while also increasing NMDAR transients and overall neuronal excitability for LTP ^47,48,54,81–84^. Together, increased PKA activity and NMDAR channel conductance elevate intracellular Ca²[ levels, leading to the nuclear translocation of ERK1/2. This ultimately leads to cAMP-response element binding protein (CREB)-dependent transcription of PRPs required for late-LTP ^85^. Consequently, PKA has been proposed to be involved in tag setting and is crucial for the early-LTP to late-LTP transition in the CA1 ^31,86^. Furthermore, persistent D1R activation promotes late-LTP in an ERK1/2-dependent manner in the CA1 ^24,87^. However, STEP suppresses LTP induction by inhibiting ERK1/2, resulting in enhanced NMDAR and AMPAR internalisation from the synaptosomal surface membrane ^52–54,88–90^. D1R activation relieves STEP-mediated inhibition on ERK1/2 via PKA, thus enabling LTP expression ^54,88,91^. Our results reveal that these mechanisms are also prevalent in the CA2. Sufficient dopaminergic and glutamatergic co-activation differentially modulate CA2 plasticity in a protein synthesis- and NMDAR-dependent manner. For the EC-CA2, this co-activation is sufficient to set tags, initiate PRP synthesis and capture them for late-LTP, albeit slowly as PKA activity gradually accumulates. For the SC-CA2, prior D1R priming enables LTP induction only upon WTET/STET. This effect involves upregulated PKA and ERK1/2 activity and downregulated STEP activity. These primed and tetanised SC-CA2 synapses capture available PRPs for LTP expression. Since silencing the EC-CA2 abolishes this effect, it suggests that the SC-CA2 is unable to recruit adequate PRPs for LTP. Thus, the SC-CA2 relies on EC-CA2-generated PRPs for induction and sustained potentiation. This way, both EC-CA2 and SC-CA2 enjoy the benefits of synaptic cooperation.

It is imperative to elucidate the effects of dopamine signalling in the CA2 as the social deficits associated with schizophrenia involve crosstalk between the hippocampus and dopaminergic sources in the brain ^92–95^. Interestingly, chlorpromazine-mediated antagonism of Dopamine-2 receptors (D2R) also results in upregulated PKA activity while attenuating STEP in the striatum ^96,97^. The interplay between STEP and PKA mediates the beneficial effects of neuroleptics for treating schizophrenia ^93,98,99^. In contrast to D1Rs, D2Rs are G_i/o_ protein-coupled receptors that exert an inhibitory effect - consequently, D2R activation in the CA1 results in impaired LTP and deficits in hippocampus-dependent learning ^100^. Given the distinct affinities of D1Rs and D2Rs for dopamine, this area presents a compelling avenue for future research ^101^. While earlier studies supported the dopamine hypothesis of schizophrenia, as the efficacy of antipsychotics like chlorpromazine depends on its affinity for D2R, they do not wholly explain reports of NMDAR antagonist-induced schizophrenia-like behaviour ^89,96,97^. Recent reports highlight the involvement of intracellular targets like PKA, STEP and ERK1/2 as essential players in mediating beneficial neuroleptic effects in the striatum ^89,99^. Since dopaminergic and glutamatergic signalling converges on STEP, it validates our observations in the hippocampal area CA2.

In the non-pathological state, the CA2 interneuron population is relatively denser than the other hippocampal subfields ^102^. However, post-mortem studies from schizophrenic patients report a great loss of parvalbumin-positive interneurons in the CA2 ^103^. The CA2 alone is capable of driving CA1 firing ^8^. If not kept in check by feedforward inhibition from the CA3, CA2 firing may lead to CA1 hyperactivity. Furthermore, dopamine and glutamate dysfunction in the hippocampus are associated with schizophrenia pathophysiology ^10,21,27,30,92,104,105^. Consequently, the loss of CA2 inhibitory interneuron population coupled with dopaminergic hyperactivity may contribute to CA1 malfunction and hippocampus-related behavioural deficits seen in schizophrenia. Despite socially rewarding experiences, schizophrenic patients have impairments in forming or updating their internal representations of social information ^106^. Consequently, they exhibit social withdrawal and reclusive behaviour ^107^. Given the crucial role of the CA2 in social memory, it is likely that dopaminergic projections that innervate the area CA2 influence social memory consolidation and subsequent social behaviour, or the lack thereof.

A limitation of *in vitro* field electrophysiology is that behavioural outcomes cannot be ascertained. Therefore, the behavioural consequences of dysregulated dopaminergic transmission in the CA2 remain uncertain. Additionally, direct dopamine agonist application and the brain slice interface chamber cannot fully recapitulate the physiological setting. As such, future studies could trace dopaminergic innervations to the CA2 via retrograde viral infection, monitor *in vivo* dopaminergic activity via G-protein-coupled receptor-activation-based dopamine (GRAB_DA_), and investigate changes in social memory-related behaviour before and after excitatory optogenetic activation. Our findings here set the groundwork for elucidating the effect of dopaminergic transmission in the area CA2 and thereafter the development of meaningful therapies for social-related neuropsychiatric disorders.

## Acknowledgements

This work was supported by NUHS seed fund (NUHSRO/2024/089/T1/Seed-Mar24/02) and Ministry of Health (MOH-000641-00) to SS, and NUS Research Scholarship to KC.

---

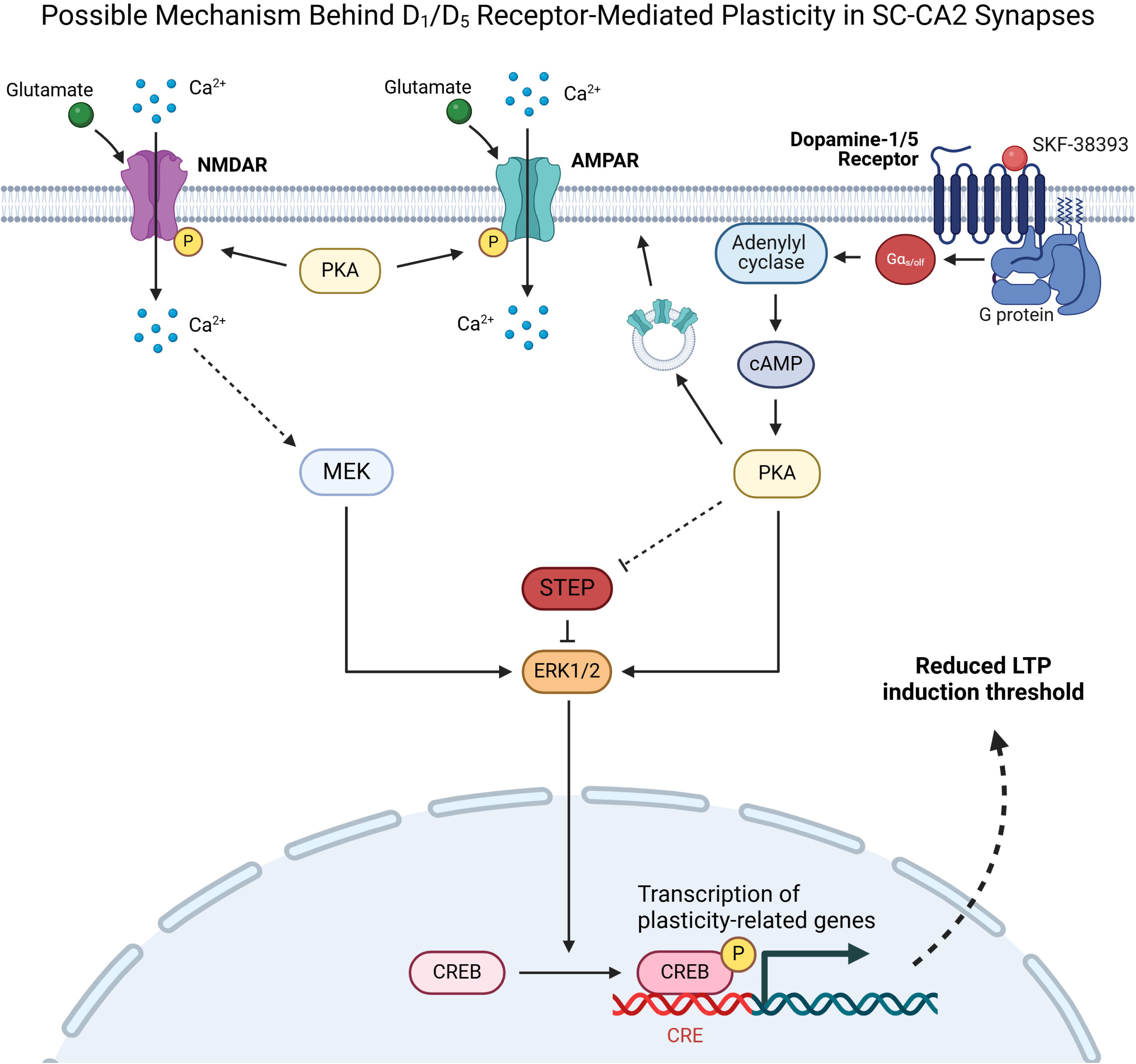

